# Unattended but actively stored: EEG dynamics reveal a dissociation between selective attention and storage in working memory

**DOI:** 10.1101/320952

**Authors:** E. Gunseli, J. Fahrenfort, D. van Moorselaar, K. Daoultzis, M. Meeter, C. N. L. Olivers

## Abstract

Selective attention plays a prominent role in prioritizing information in working memory (WM), improving performance for attended representations. However, it remains unclear what the consequences of selection are for the maintenance of unattended WM representations, and whether this results in information loss. Here we tested the hypothesis that within WM, selectively attending to an item and the decision to stop storing other items involve independent mechanisms. We recorded EEG while participants performed a WM recall task in which the item most likely to be tested was cued retrospectively. By manipulating retro-cue reliability (i.e. the ratio of valid to invalid cue trials) we varied the incentive to retain uncued items. Contralateral alpha power suppression, a proxy for attention, indicated that, initially, the cued item was attended equally following high and low reliability cues, but attention was sustained throughout the delay period only after high reliability cues. Furthermore, contralateral delay activity (CDA), a proxy for storage, indicated that non-cued items were dropped sooner from WM after highly reliability cues than after cues with low reliability. These results show that attention and storage in WM are distinct processes that can behave differently depending on the relative importance of WM representations, as expressed in dissociable EEG signals.

## Introduction

Working memory (WM) is essential to storing and manipulating information online for a variety of cognitive tasks^1–4^. However, its capacity is limited^5,6^ and thus only the most task-relevant information should be selected for storage in WM^7,8^. Attention is the mechanism by which task-relevant representations are prioritized and there is now a large body of evidence showing that attention and WM are heavily intertwined^9–12^, such that attention may be crucial to successfully maintain an item in WM^13–21^. However, more recently alternative theoretical frameworks have been proposed that argue that storage of an item in WM should be dissociated from prioritization of (i.e. attending to) that item^22–27^. Thus, there is no consensus yet on the relationship between WM and attention.

Much of the evidence for a central role of selective attention in WM storage comes from studies using retrospective cues. Such “retro-cues” are presented after the to-be-remembered items have been taken away and indicate which of the memory representations is most likely to be tested and thus is the most task-relevant. Because retro-cues are presented after memory encoding, they act on stored WM representations rather than on encoding of stimuli. Nevertheless, retro-cues have been suggested to result in the attentional selection of the cued representation within WM in a similar way as attentional selection operates during perception, relying on highly overlapping neural networks^28–30^. This selection in turn has been claimed to improve storage and/or increase the accessibility of the cued item within WM^31–33^. The behavioral consequence is better memory performance for the attended representation compared to a ‘no-cue’ or neutral condition where all items are presumably equally attended^34–36^.

The finding that retrospectively cueing attention to a representation improves memory performance does not in itself prove that attention plays a necessary role in the maintenance of that representation. For that, it is necessary to show that unattended items actually suffer from attention being cued elsewhere, relative to when attention is directed equally to all items. However, so far, the fate of *unattended* WM representations has been unclear. Memory performance for unattended representations can be tested by probing a non-cued representation on a minority of the trials. A lower memory performance on these invalid cue trials compared to neutral or no-cue trials is referred to as an ‘invalid cueing cost’. Such invalid cueing costs have indeed been found in some studies, and have been taken as evidence that attention is necessary for WM storage^37–39^. However, using very similar cueing procedures, a number of other studies did not find such invalid cueing costs^21,23,30^, suggesting a dissociation between storage and selection.

As we have proposed earlier^41^, the fate of non-cued items might depend on their perceived future relevance, as inferred from the reliability of the retro-cue (i.e. the proportion of valid to invalid retro-cue trials). Typically, studies that failed to observe invalid cueing costs used lower retro-cue reliabilities^31,40,42^ than studies that observed invalid cueing costs^32,37,39,43^. In a behavioral study, we observed invalid cueing costs only when the retro-cue had a high reliability (i.e., 80% valid), but not when it had a lower, but still above-chance reliability (i.e., 50% valid, with chance level being at 25% in both conditions). While the presence of invalid retro-cue costs varied with retro-cue reliability, benefits of *valid* retro-cues were present in both conditions, though they were larger for 80% valid cues.^41^ This can explain the discrepant findings in the literature if we assume that attending to an item in WM can be dissociated from the decision to either continue or cease storage of remaining items. For both moderately and highly reliable cues it is beneficial to attend to an item, as it is more likely to be tested than uncued items. However, only for highly reliable cues it is also worth dropping the uncued items from memory, while for moderately reliable cues it is actually worth holding on to the uncued items.

Although our behavioral work provides initial evidence for the idea that attention to and storage of an item should be dissociated when interpreting the effects of retro-cueing, there is an alternative scenario that can explain the reliability effects on performance for uncued items, in which increasing the retro-cue reliability results in more attention to the selected representation without affecting the probability with which the unattended representations are dropped from WM. Under this scenario, items in principle remain stored in WM regardless of cue reliability, but they become more vulnerable to interference from the test display when unattended. The test display is in itself a stimulus that may overwrite a fragile memory representation, and it has been proposed that attention protects against such interference^44–47^. This would then result in larger invalid cueing costs for highly reliable cues, even if unattended items were still stored until the test display. While a differential storage account predicts that the decision to drop an uncued item is made during the retention interval, the protection against interference account predicts that nothing happens to uncued items during the retention interval and that performance differences result from processes during test. Because behavioral methods only measure the final outcome, they are blind to the underlying mechanisms during retention and therefore cannot differentiate between these scenarios.

To more directly investigate if and how retro-cue reliability affects attention and storage in WM prior to the test, we used EEG recordings to measure these processes in a time-resolved manner during the retention interval. The experimental procedure is shown in Figure 1A. We used a continuous report WM task to obtain a sensitive measure of memory performance. The memory display contained three line segments of different orientations, one on the vertical midline and the other two presented left and right from fixation. After a blank interval, a retro-cue indicated which of the memory representations was most likely to be tested by retrospectively pointing to its location in the memory display. Only lateral cue trials were included for the EEG analysis since both of our EEG indices of interest (see below) required a lateral asymmetry in the location of the attended and stored item. Critically, to vary the incentive to also retain the uncued items, we manipulated the retro-cue reliability (i.e. the proportion of valid to invalid trials) across blocks: The cue was 50% valid in half the number of blocks, and 80% valid in the other half.

**Figure 1.**
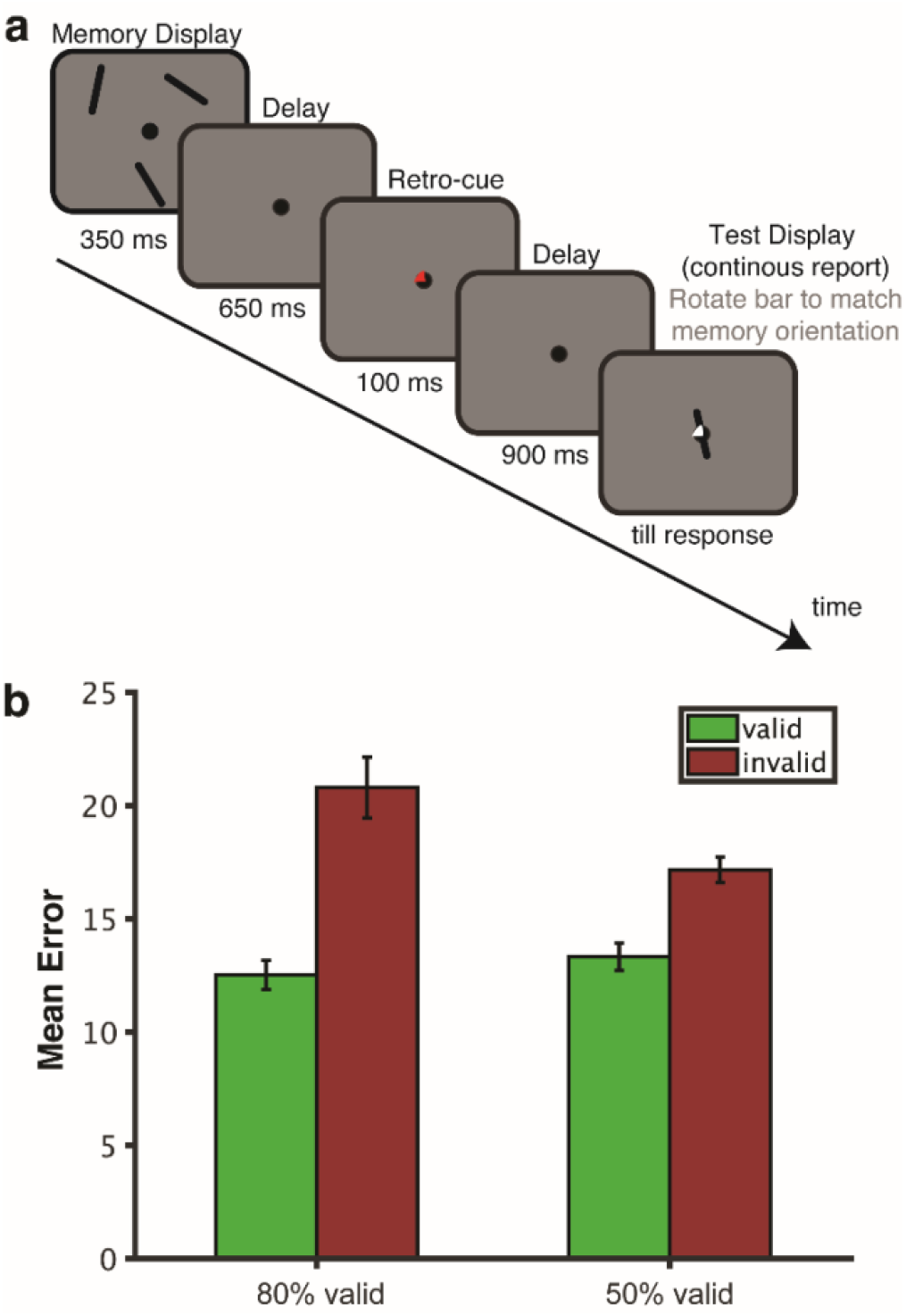
**(A)** The retro-cue experimental procedure. Participants were asked to remember the three orientations shown in the memory display. After a blank interval, a retro-cue was presented pointing to the location of the item (in this example top-left) that was most likely to be tested. Retro-cues were not always valid. Following a second blank interval the test display was presented during which participants were asked to rotate a randomly-oriented bar to match the orientation of the tested item (which in this example is the item presented on top-left, hence the retro-cue was valid). **(B)** Average error for reporting the probed orientation in each condition. The valid and invalid trials are shown in green and red respectively. Error bars represent standard errors of the mean for normalized data, i.e. corrected for between-subjects variance (Cousineau, 2005). Retro-cue validity effect was larger for highly reliable cues than less reliable cues.

As a proxy for *attention* being directed within WM we used contralateral power suppression in the alpha band (8-14 Hz). Alpha power over the parietal-occipital electrodes on the hemisphere contralateral to the attended item has been found to be reduced relative to the ipsilateral electrodes, both during perception and during post-perception within WM^48–53^. We hypothesized that if the cued item is attended more during storage for highly reliable cues, then we should observe a larger contralateral alpha suppression for highly reliable retro-cues than for cues of lower reliability. As a marker for *storage* we focused on the CDA, which is a sustained negativity over the parietal-occipital electrodes on the hemisphere contralateral to remembered stimuli. It has been observed to be sensitive to the visual WM load, and converging evidence suggests that it is an index of visual WM storage^54–57^. We reasoned that if non-cued representations are dropped following a retro-cue, then a CDA should emerge contralateral to the attended item, since dropping an item on one side results in an imbalance in the number of items stored in each hemisphere^58^. If, as we hypothesized, the likelihood of dropping an item depends on retro-cue reliability, we should see a stronger CDA emerge in the high reliability condition than in the low-reliability condition. Alternatively, if retro-cue reliability has no effect on storage, we should see no differential CDA, and only find attentional effects as expressed through alpha suppression.

## Method

Thirty-two healthy volunteer university students (ages 18-35) participated in the experiment for course credit or monetary compensation. Two participants were excluded; one due to excessive noise in their EEG recordings and one due to poor behavioral performance (see Data Analyses), leaving 30 participants (22 female) of whom the data was analyzed. The study was conducted in accordance with the Declaration of Helsinki and was approved by the Scientific and Ethical Review Board (Dutch abbreviation: VCWE) of the Faculty of Behavioral and Movement Sciences. Written informed consent was obtained prior to the experiment. Data sets are available online on Open Science Framework at https://osf.io/bgpxc/?view_only=3b8dd8f9e4fa42d68ac84db90f76e25d

The procedure is shown in Figure 1. Each trial started with the presentation of the fixation circle of radius 0.33°, for a duration jittered between 1200-1600 ms. Then, the memory display was presented for 350 ms. It consisted of three black oriented bars (2.08° x 0.25° visual angle) located at 60 (top right), 180 (bottom) and 300 (top left) degrees relative to the top of an imaginary circle of radius 3.50°. We used a memory load of three items in order to tax WM without contaminating measurements with non-encoded items. The orientation of each bar was chosen at random with the restriction that bars within the same trial differed by at least 10°. The retro-cue was presented for 100 ms following a blank interval of 650 ms during which only the fixation circle was presented. The retro-cue was identical to the fixation circle except that one quarter (90°) was now filled with either red, 27.08 Cd/m^2^, or green, 24.10 Cd/m^2^, depending on the reliability condition (order counterbalanced). For the initial practice phase where the cue was 100% valid, the retro-cue fill color was orange (53.46 Cd/m^2^). Following the retro-cue, there was a blank interval of 900 ms in which only the fixation circle was presented. Then the test display was presented till response. It contained a probe cue pointing to the location of the tested representation and a randomly oriented probe bar that were both presented at the center of the screen. This probe cue was the same as the retro-cue except that the filling color was white. Participants were asked to indicate the orientation of the bar at the tested location as precise as possible by rotating the probe bar using the mouse and pressing the left mouse button. After a mouse response was made, the correct orientation was indicated by a central white bar for 100 ms. The screen was empty during the inter-trial interval which was jittered between 1200-1600 ms.

The retro-cue was 80% valid for half of the experiment and 50% valid for the other half (order counterbalanced). There were 10 blocks of 50 trials. Each validity condition (i.e. valid and invalid) was randomly intermixed within each block. Before each reliability condition, participants were informed about the validity ratio of the retro-cue and they performed a practice session of 25 trials to get used to this particular validity ratio. Moreover, at the beginning of the experiment, there was an initial practice session of 25 trials with a 100% valid cue to make participants familiar with using the experimental procedure. At the end of each block, participants received feedback on block average and grand average error (i.e. the difference between the original tested orientation and the responded orientation).

### EEG Data Acquisition

The electroencephalogram (EEG) and electro-oculogram (EOG) were recorded from 70 sintered –AG/AgCl electrodes positioned at 64 standard International 10/20 System sites and 6 external locations mentioned below, using the Biosemi ActiveTwo system (Biosemi, Amsterdam, the Netherlands). We did not perform impedance measurements as recommended by Biosemi: High impedance has been suggested to have minor impact on data quality in cool and dry environments^59^. The vertical EOG (VEOG) was recorded from electrodes located 2 cm above and below the right eye, and the horizontal EOG (HEOG) was recorded from electrodes 1 cm lateral to the external canthi. The VEOG was used in the detection of blink artifacts, and the HEOG was used in the detection of horizontal eye movement artifacts. Electrophysiological signals were digitized at 512 Hz.

### Data Analysis

#### Behavior

Error scores on the memory test were calculated as the difference between the original orientation of the tested memory bar and the orientation of the response. One participant with an average absolute error value higher than 2.5 standard deviation above the grand average of the group was excluded from analysis. Absolute error for the tested item was entered into a repeated-measures ANOVAs with the within-subjects factors of retro-cue reliability (80% valid; 50% valid) and retro-cue validity (valid; invalid).

We also estimated the guess rate (i.e. reporting a random orientation), swap rate (i.e. reporting a non-tested representation), and sd (inverse of precision) based on the width of the response distribution around the target using MemToolbox (memtoolbox.org)^60^. To test if the large trial number imbalance between valid and invalid trials in 80% valid condition (200 vs 50) has an impact of parameter estimates, we took 200 trials in 80% valid condition and downsampled 50, 100, and 150 trials over 20 iterations. We found that having a lower number of trials significantly inflates guess rate and swap rate estimates, but more importantly increases the variability of each estimate (i.e. decreases their reliability). Thus, we report only raw errors and not the parameter estimates from the swap model.

#### EEG analysis: General

Only lateral cue trials were included for the EEG analysis since both of our EEG indices of interest require a lateral asymmetry in the location of the attended or stored item. All EEG analyses were carried out using the EEGLAB toolbox^61^ and custom scripts implemented in MATLAB (The MathWorks, Inc., Natick, MA). Due to unknown reasons, there were three participants who had parts of EEG data missing (10, 11 and 26 trials). Noisy electrodes were interpolated using the “eeg_interp.m” function of EEGLAB with the spherical interpolation method, which resulted in the interpolation of three electrodes each for two participants (FC2, C6, PO3; CP3, PO3, P4). None of these electrodes were used in the statistical analysis. Trials with recording artifacts (muscle noise and slow drifts) and ocular artifacts (blinks and eye movements) were rejected manually by visually inspecting the EEG and EOG electrodes respectively. Artifact rejection was performed in the absence of any knowledge about the conditions. Individuals were excluded from analyses if, after all the artifact rejections, the remaining number of trials per condition was lower than 80 trials. This led to the rejection of one participant. For the remaining participants, on average 9.8% of all trials were rejected due to artifacts, leaving on average 139 and 141 lateral cue trials for analysis (with a minimum of 105 and 111 trials), for 80% valid and 50% valid blocks respectively.

#### ERP analysis: CDA

ERPs were computed with respect to a 200 ms pre-stimulus baseline period, between −500 to 1500 ms around the retro-cue display and were re-referenced offline to the average of left and right mastoids. The data was filtered with an IIR Butterworth filter with a bandpass of 0.01 – 6 Hz. We chose an upper limit of 6 Hz to remove alpha-band activity from the CDA calculation to fully isolate the two signals. However, our main findings (i.e. CDA being significant only for 80% valid condition early in the trial, but for both 50% and 80% valid conditions late in the trial) were identical when the CDA was calculated using a bandpass filter of 0.01 – 40 Hz. Signal was resampled at 500 Hz using “pop_resample.m” function of EEGLAB.

The CDA was calculated as the difference waves between electrode sites contralateral versus ipsilateral to the location of the retro-cued item. Previous studies measuring the CDA have typically found maximal values at posterior/occipital electrodes and started measuring the CDA at ~300-400 ms from the onset of the memory display following the N2pc (~200-300 ms) which signals individuation of selected items^54,62,63^. Based on these studies and visual inspection of the topographic distribution of lateralized voltage in posterior/occipital regions we calculated the CDA at P7/8, PO7/8, and O1/2 as contralateral minus ipsilateral to the retro-cud location starting from 400 ms following the retro-cue onset till the onset of the test display (i.e. 900 ms after cue onset). The CDA averaged across electrode pairs and times of interest was entered into a repeated-measures ANOVA with the within-subjects factor of retro-cue reliability (80% valid; 50% valid). Average CDA values were also tested against zero using one-sample t-tests. Additionally, given that CDA is a sustained negativity at electrodes contralateral to items stored in WM, we hypothesized that if the reliability effect on CDA reflects a boost for the representation of the cued item, then it should be evident in the signal contralateral to the *cued* item, while if it reflects dropping of the non-cued item then the reliability effect should be observed in the signal contralateral to the *non-cued* item (i.e. ipsilateral to the cued item). To test this, the average contralateral and ipsilateral signals were entered into a repeated-measures ANOVA with within-subjects factors of laterality (contralateral; ipsilateral to the cued item) and retro-cue reliability (80% valid; 50% valid).

In order to investigate the dynamic time course of the reliability effect, the CDA at each time point for each reliability condition were tested against chance and also against each other at a group level using cluster-based permutation testing by estimating the permutation *p*-value using a Monte Carlo randomization procedure^64^. For this analysis, we randomly shuffled the condition labels (e.g. 50% valid vs. 80% valid) 1000 times to approximate the null distribution of the t statistic. The *p*-value was the proportion of iterations out of 1000 where the absolute randomly shuffled condition difference was larger than the absolute actual condition difference (two-tailed; thus 0.025 for positive and 0.025 for negative). Multiple comparisons correction was established using cluster-based permutation testing^64^. First, four or more temporally adjacent data points with a p-value smaller than 0.05 were clustered together (again two-tailed). Then, a cluster-level statistic was calculated by taking the sum of the t-values within each cluster, separately for positive and negative clusters. The p-value for each cluster was calculated as the number of times the sum of the absolute t-values within the cluster under random permutation exceeds that of the t-values within the observed cluster. A cluster was considered significant if the calculated p-value was smaller than 0.05. Note that, there are two separate t-tests described above as part of cluster-based permutation testing. First to determine time points that form a cluster, second to determine significant clusters.

#### Alpha band (8-14 Hz) power

Alpha band (8-14 Hz) power analysis was performed using the same trials as in the CDA analysis. Prior to the calculation of power, the signal was epoched between −1000 to 2000 ms around the onset of the cue display. We chose a larger window compared to the CDA analysis in order to avoid contaminating the results from edge artifacts that result from applying a band-pass filter at the edges of an epoch^64^. To isolate alpha-band activity, we bandpass-filtered the raw EEG between 8 and 14 Hz using a two-way IIR Butterworth filter of 4-th order as implemented by the ft_preproc_bandpassfilter.m function of FieldTrip Toolbox^66^. Then, in order to produce a complex analytic signal, 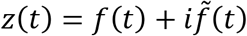, we applied a Hilbert transform (MATLAB Signal Processing Toolbox) to the band-pass filtered data, *f*(*t*), where 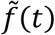 is the Hilbert Transform of *f*(*t*) and 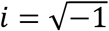, using the following Matlab syntax:

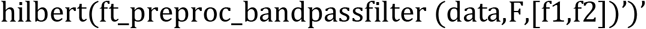

where data is the raw EEG (trials x samples), F is the sampling frequency (500 Hz), f1 and f2 are the boundaries of the frequency band to be isolated (8 Hz and 12 Hz respectively). We computed instantaneous power by taking square of the complex magnitude of the complex analytic signal. After calculating the power, the epochs were reduced to −500 to 1500 around the retro-cue display. Power data was baseline normalized separately for each condition (i.e. 50% valid left-cue, 50% valid right-cue; 80% valid left-cue; 80% valid right-cue) with decibel (dB) conversion, using −400 to −100 ms relative to the retro-cue onset as baseline^65^. The dB normalized data was averaged separately for contralateral and ipsilateral in respect to the side of the cued item at the electrode pairs of interest (P7/P8, PO7/PO8, O1/O2). Contralateral alpha suppression was calculated as the difference between the contralateral and ipsilateral dB normalized power values.

Contralateral alpha band power averaged across electrodes and times of interest (400 – 900 ms, which is chosen to be the same time interval as the CDA analysis) were compared between 50% and 80% valid conditions using a repeated measures ANOVA. Average contralateral alpha band power values were also tested against zero using one-sample t-tests. Since we performed a reliability analysis separately for contralateral and ipsilateral signal for the CDA, for completeness we also performed it for lateral alpha power. Power values contralateral and ipsilateral power relative to the direction of the cued item were averaged across the time window of interest entered into a repeated-measures ANOVA with within-subjects factors of laterality (contralateral; ipsilateral to the cued item) and retro-cue reliability (80% valid; 50% valid). Lastly, contralateral power suppression at each time point for each reliability condition were tested against chance and against each other at a group level with the same cluster-based permutation test as in the CDA analysis.

#### Investigating the effects of eye movements on EEG measures of interest

Eye movements can contaminate EEG signal as a result of a potential difference between the cornea and fundus of the eye^67^. Therefore, even though we rejected trials with eye movements as reflected in HEOG electrodes, it is possible that subtle differences in the number of eye movements across retro-cue reliability conditions might have spuriously produced differences in CDA and lateralized alpha-band power. In order to evaluate this possibility, first we calculated the average potential difference between left and right HEOG electrodes during our time window of interest (i.e. 400 – 900 ms) separately for trials where the cued item was on the left vs. the right hemifield and then averaged their absolute values. This allowed us to assess the amount of horizontal eye movements, as previous work has shown that an HEOG voltage difference of 3.2 μV corresponds to 1 visual degrees of horizontal saccades. Second, we compared this measure across 80% valid and 50% valid conditions using a repeated measures ANOVA to test if the amount of eye movements differed across reliability conditions. Third, to evaluate whether trial-by-trial differences in the amount of eye movements might have affected our EEG measures of interest to a different degree across reliability conditions, we calculated Pearson’s correlation coefficients across trials, per participant, per condition between our measures of interest (i.e. CDA and contralateral alpha suppression) and contralateral minus ipsilateral HEOG averaged across our time-window of interest (i.e. 400 – 900 ms). Then, Pearson’s correlation coefficients per participant and per condition were Fisher transformed to ensure normality and were entered into a repeated measures ANOVA with the factor of retro-cue reliability. Also, given that CDA and contralateral alpha suppression effects were specific to early and late time intervals respectively, we repeated this analysis separately for the time windows at which cluster-based permutation tests showed a significant reliability effect on CDA (400 - 514 ms) and contralateral alpha suppression (724 - 900 ms). For these trial-level correlation analyses, we calculate alpha-band power per trial by using a single-trial baseline normalization approach instead of using a common baseline per condition that is averaged across trials.

#### Correlation between EEG measures and behavior

To test if differences across reliability conditions in our EEG measures of interest predict behavior, we calculated the difference in invalid cueing cost (i.e. error in invalid trials minus error in valid trials) between 80% valid and 50% valid conditions, and correlated this with the CDA and contralateral alpha suppression differences between these two conditions. We calculated average CDA and contralateral alpha suppression at the time window in which the condition difference was significant within the retention interval as revealed by the aforementioned cluster-based permutation tests (400-514 ms for the CDA and 724-900 ms the contralateral alpha suppression).

To test if variability in our EEG measures of interest predicts behavior, we calculated, per cell (participant; reliability condition; validity condition), Pearson’s correlations between trial-level CDA and error, and between trial-level contralateral alpha power and error (except invalid trials of 80% reliable condition; as there were too few trials in this condition). Then, average Pearson’s correlation coefficients were Fisher’s transformed to ensure normality, and were tested against zero using one-sample t-tests.

#### Correlation between CDA and contralateral alpha suppression

In order to test whether selective attention within WM predicts storage in WM on a participant level, we performed Pearson correlations between the average CDA and the average contralateral alpha suppression across participants, separately for 50% valid and 80% valid blocks. In order to investigate the same relationship at a within-participant level, we calculated Pearson’s correlation coefficients across trials, per participant, per condition between our measures of interest (i.e. CDA and contralateral alpha suppression) averaged across the time-window of interest (i.e. 400 - 900 ms). Then, Pearson’s correlation coefficients per participant and per condition were Fisher transformed to ensure normality and were tested against zero separately for 50% valid and 80% valid blocks using one-sample t-tests. Given that the CDA and contralateral alpha suppression effects were specific to early and late time intervals respectively, we also performed this analysis separately for the time windows at which cluster-based permutation tests showed significant reliability effects on CDA (400 - 514 ms) and contralateral alpha suppression (724 - 900 ms).

## Results

### Behavior

Figure 1B shows mean absolute error (i.e. the absolute difference between the actual probed orientation and the responded orientation) for each condition. There was a main effect of validity on error, *F*(1, 29) = 27.75, *p* <.001, η_p_^2^= 0.49. Errors were larger on invalid (*Mean* = 18.97, 95% CI [16.09, 21.84]) compared to valid cue trials (*Mean* = 12.92, 95% CI [11.62, 14.22]). Importantly, this validity effect (i.e. error on invalid trials minus the error on valid trials) was larger when the cue was 80% valid, *Mean* = 8.27, 95% CI [4.39, 12.15], compared to when it was 50% valid, *Mean* = 3.83, 95% CI [2.31, 5.35], as indicated by a validity x reliability interaction, *F*(1, 29) = 6.49, *p* = 0.016, η_p_^2^= 0.18. There was no main effect of reliability on error, *F*(1, 29) = 2.41, *p* = 0.131, η_p_^2^= 0.07. In sum, and as we expected, the effect of retro-cues on recall performance was larger for cues that were more reliable.

### Electrophysiology

#### Selective attention was allocated to the cued item independent of retro-cue reliability, but was sustained only for highly reliable cues

Figure 2A shows the contralateral alpha band (8-14 Hz) power suppression (i.e. the difference between the contralateral and ipsilateral power) with respect to the position of the cued item averaged across the electrode pairs of interest (P7/8, PO7/8 and O1/2). Cluster-based permutation tests showed significant contralateral alpha suppression in both the 80% valid condition (significant clusters: 362-544 ms, *t*(29) = −2.59, *p* = 0.015; and 682-850 ms, *t*(29) = −2.40, *p* = 0.011) and the 50% valid condition (significant clusters: 416-632 ms, *t*(29) = −2.74, *p* = 0.010), with initially no difference between the validity conditions. After about 600 ms however, the contralateral alpha suppression in 50% valid condition dropped back to the baseline, resulting in a significant difference between the 50% valid and 80% valid conditions in the later part of the delay period (significant cluster: 724-970 ms; *t*(29) = −2.55, *p* = 0.016). As seen in Figure 3A the retro-cue reliability affected the signal on the contralateral hemisphere relative to the cued item, *F*(1, 29) = 4.70, *p*=.038, η_p_^2^= 0.14, 95% CI [0.02, 0.56], but not the ipsilateral hemisphere, *F*(1, 29) = 0.02, *p*=.885, η_p_^2^< 0.01, 95% CI [−0.20, 0.17]. This was reflected in a significant laterality x reliability interaction, *F*(1, 29) = 8.42, *p*=.007, η_p_^2^= 0.22. These results suggest that early in the trial the cued item was attended independent of retro-cue reliability, but that attentional prioritization was sustained for a longer period of time when cue was more reliable.

**Figure 2.**
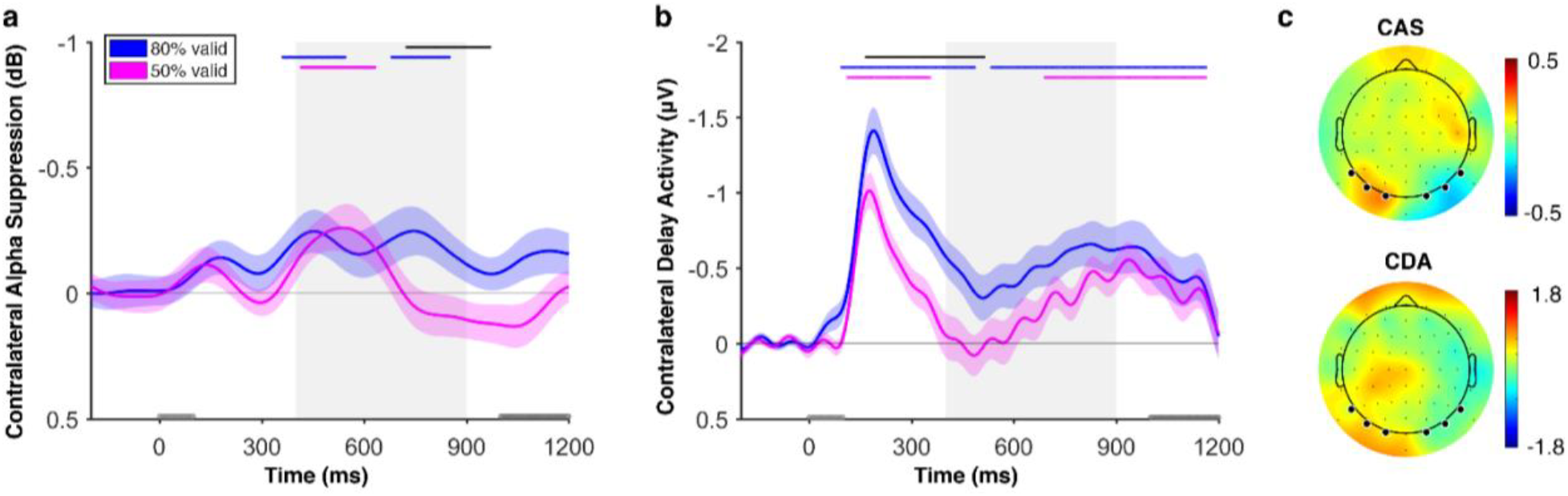
**(A)** Contralateral alpha suppression, and **(B)** CDA as indices of selective attention and storage in WM respectively, both time-locked to the onset of the retro-cue, are shown in different colors for 80% valid and 50% valid conditions. The gray area shows the time window of interest (400-900 ms). The gray rectangles on the x-axis show the timing of the retro-cue (0-100 ms) and the test display (from 1000 ms till response, which extends till 1200 ms on the plots). Markers along the top of each plot indicate the time points at which either the difference between the EEG measures in 80% valid and 50 % valid conditions (black) or the EEG measure itself for each condition (blue for 80% valid and magenta for 50% valid) were significantly different than zero as determined by a cluster-based permutation test (*p*<0.05; two-tailed). For highly reliable cues, the cued item was attended and non-cued items were dropped from WM. For less reliable cues, non-cued items were initially unattended but were kept in WM until about the onset of the test display. Error bars represent standard error of the mean. (C) The scalp maps of contralateral alpha suppression (CAS) and contralateral delay activity (CDA) averaged across the time window of interest (400-900 ms after the cue onset) calculated as the difference of trials when the cued item was on the left minus when it was on the right hemifield collapsed across 80% valid and 50% valid conditions. The dots on the scalp map shows the positions of the EEG electrodes. The thicker dots on the posterior side of the scalp map shows the electrodes used for calculating CAS and CDA.

**Figure 3.**
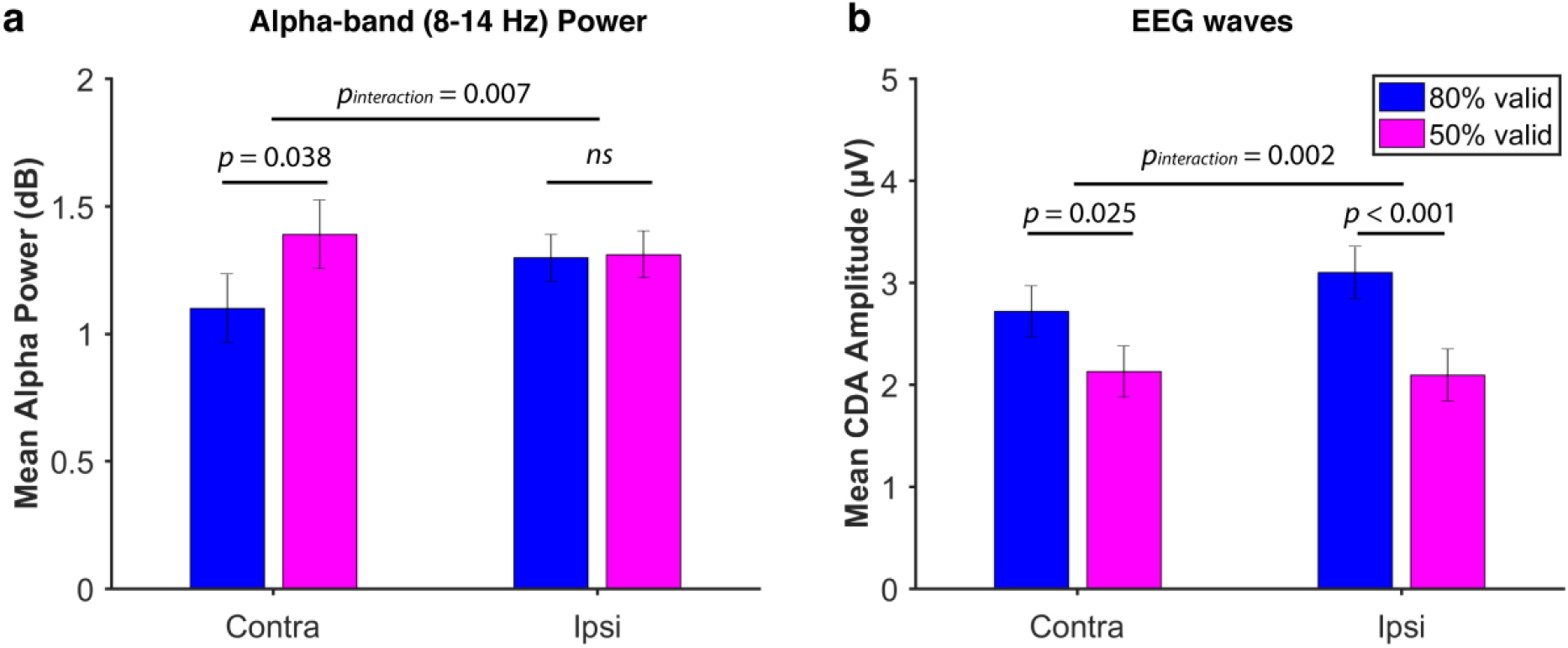
**(A)** Alpha power and **(B)** EEG waves contralateral and ipsilateral to the side of the cued item averaged across the electrodes of interest during the time window in which cluster-based permutation testing revealed a significant reliability effect during the retention interval (i.e. 724-900 ms for alpha-band power and 400-514 for the CDA). 80% valid and 50% valid conditions are shown in different colors. The reliability effect was stronger at electrodes contralateral to the *non-cued* item for CDA, but contralateral to the *cued* item for alpha power. This is in line with the claim that the contralateral alpha suppression reflects attentional selection of the cued item while the CDA reflects dropping of the non-cued item. The error bars represent the standard error of the mean for the difference between 80% valid and 50% valid conditions separately for contra and ipsilateral values.

#### Non-cued items were dropped from WM earlier and more often after highly reliable cues

Figure 2B shows the difference between contralateral and ipsilateral waveforms in respect to the location of the retro-cued item, averaged across the electrode pairs of interest (P7/8, PO7/8 and O1/2). Cluster-based permutation tests showed that for 80% valid blocks there was a significant CDA both during early and late parts of the retention interval (significant clusters: 94-486 ms, t(29) = −6.06, p < 0.001, and 534-1164 ms, t(29) = −3.44, p = 0.002, after the retro-cue onset), while for 50% valid blocks there was a CDA only later in the trial, just before the test display (significant cluster: 690-900 ms, t(29) = −3.42, p = 0.002). Interestingly, and in contrast to the alpha suppression here the difference between 80% valid and 50% valid conditions was mainly apparent early in the delay period (166-514 ms, t(29) = −3.31, p = 0.002), rather than late. To compare the effect of retro-cue reliability on contra and ipsilateral signals separately we averaged the EEG voltage across the time interval at which the condition difference was significant as revealed by cluster-based permutation test. As seen in Figure 3B, the effect of retro-cue reliability was larger on the ipsilateral hemisphere, *F*(1, 29) = 5.55, *p* = 0.025, η_p_^2^= 0.16, 95% CI [0.08, 1.10], compared to the contralateral hemisphere, *F*(1, 29) = 15.52, *p* < 0.001, η_p_^2^= 0.35, 95% CI [0.48, 1.52], relative to the cued item. This difference was reflected in a significant laterality x reliability interaction, *F*(1, 29) = 11.24, *p*=.002, η_p_^2^= 0.28. This result is in contrast with the findings in typical CDA studies where participants store items presented in a single hemifield and the memory load mainly affects the signal contralateral to the memory items^68^. This dissociation in line with our conclusion that the CDA here reflects dropping of the non-cued item instead of attending the cued item. In sum, the CDA results suggest that non-cued items were dropped from WM right after the retro-cue after highly reliable cues but only later in the retention interval after less reliable cues.

#### Correlations between EEG and Behavior

To test if differences across reliability conditions in our EEG measures of interest predict behavior, we calculated the difference in invalid cueing cost (i.e. error in invalid trials minus error in valid trials) between 80% valid and 50% valid conditions, and correlated this with the CDA and contralateral alpha suppression differences between these two conditions. We calculated average CDA and contralateral alpha suppression at the time window in which the condition difference was significant within the retention interval as revealed by the aforementioned cluster-based permutation tests (400-514 ms for the CDA and 724-900 ms the contralateral alpha suppression). There was no correlation between contralateral alpha suppression and behavior (*R^2^* < 0.01, *p* = 0.92). However, as seen in Figure 4, there was a significant correlation between CDA and behavior: Participants who had a larger invalidity cost in 80% valid condition compared to 50% valid condition also had a larger CDA difference between these reliability conditions, *R^2^* = 0.23, *p* = 0.007. Note that although in the expected direction, with N=30, and four participants having invalidity costs rather far from the mean, this correlation should be treated with some caution.

**Figure 4.**
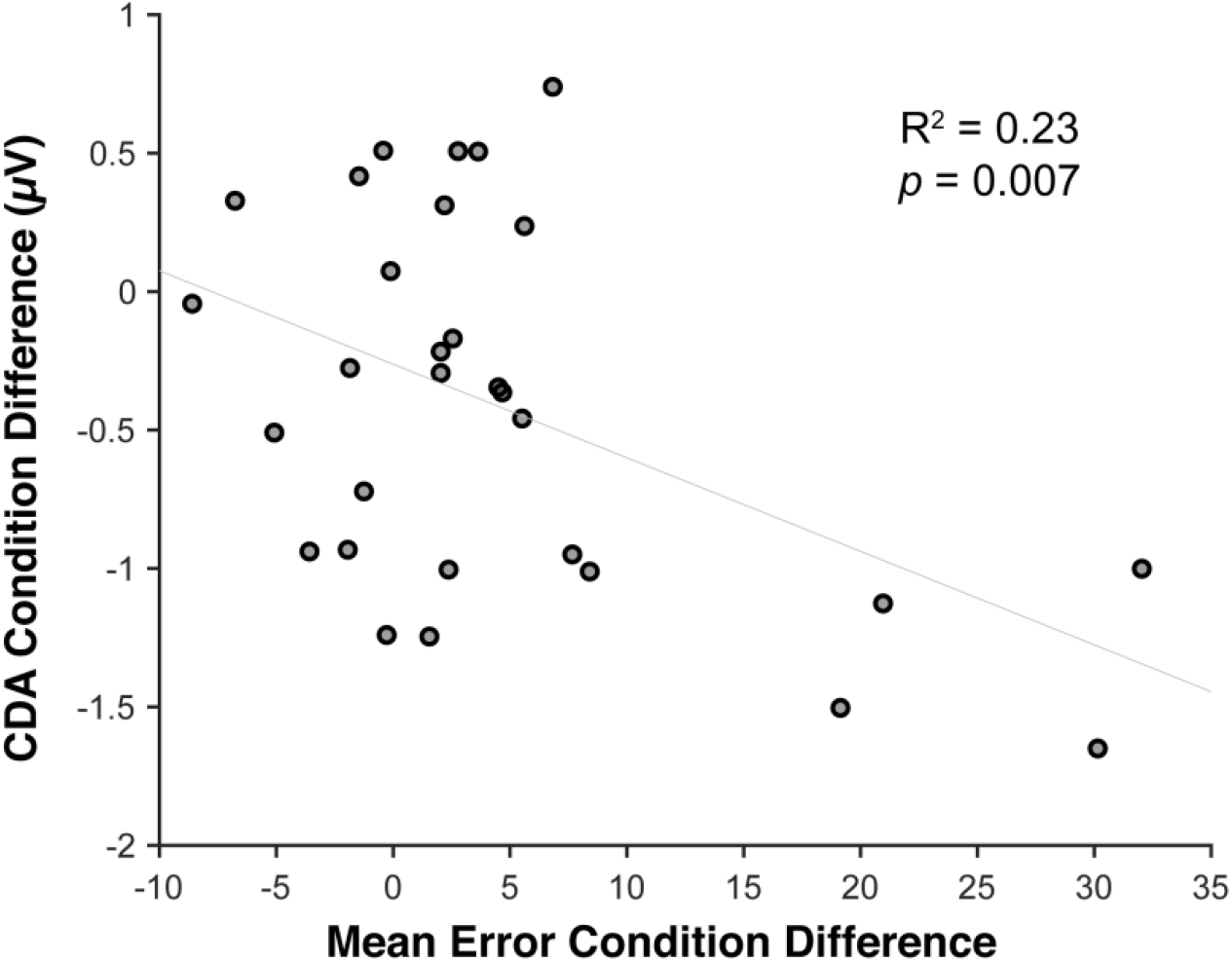
Correlation between reliability effects on CDA and invalid cueing cost. The x-axis shows the invalid cueing cost difference between reliability conditions and the y-axis shows the CDA amplitude difference between reliability conditions. A larger CDA difference is correlated with a larger invalid cueing cost on behavior.

In addition, to test if variability in our EEG measures of interest predicts behavior, we computed at the level of individual data cells (participant; reliability condition; validity condition), the Pearson’s correlations between trial-level CDA and error, and between trial-level contralateral alpha power and error (except invalid trials of 80% reliable condition; as there were too few trials in this condition). There was a weak but reliable correlation between contralateral alpha power and error for 50% reliable valid trials, mean *R^2^* = 0.01, *t*(29) = 2.28, *p* = 0.029. Larger (i.e. more negative) contralateral alpha suppression predicted smaller errors. No other correlation was significant (*p*s>0.21).

#### Correlations between CDA and Contralateral Alpha Suppression

To further test whether selective attention within WM relates to storage, we ran both across-participants and within-participants (i.e. across trials) correlation analyses between CDA and contralateral alpha suppression. There was no significant correlation between average CDA and average contralateral alpha suppression across participants for 80% valid blocks (*R^2^* = 0.01, *p* = 0.54) or 50% valid blocks (*R^2^* = 0.12, *p* = 0.86). Separate correlation analyses for early and late time windows also failed to reveal any correlation between the CDA and the contralateral alpha suppression (*R^2^* < 0.03, *p*s > 0.342). On a within-participants level, average Fisher transformed Pearson’s correlations between CDA and contralateral alpha suppression were no different from zero for 80% valid blocks, *t*(29) = 1.75, *p* = 0.090, 95% CI [−0.004, 0.054], or for 50% valid blocks, *t*(29) = −1.02, *p* = 0.315, 95% CI [−0.06, 0.02]. This was also the case for early and late time windows (all *p*s > 0.211). Together, these results suggest that selective attention to one item (as indexed by alpha suppression) did not predict storage of the other items (as indexed by the CDA) in WM on a trial level or on a participant level.

#### Ruling out the effects of eye position on EEG measures

In order to eliminate the effects of eye movements on EEG signal we removed trials with eye movements from analyses. However, subtle but systematic eye movements toward the cued item can generate electrical potentials that can impact the EEG signal significantly: If the amount or magnitude of eye movements differ across reliability conditions, this can account for the differences in CDA and contralateral alpha suppression differences we observed between these two conditions. In order to evaluate this possibility, we calculated the difference in voltage between left and right HEOG electrodes and averaged it across the time window of interest (400-900 ms after the cue onset). The average HEOG value was 1.41 μV, which corresponds to saccades of less than only 0.09 visual degrees.^67^ Moreover, average HEOG value was not significantly different between 80% valid blocks (*M* = 1.57 μV, *SEM* = 0.20) and 50% valid blocks (*M* = 1.26 μV, *SEM* = 0.21), *F*(1, 29) = 1.39, *p* = 0.248, η_p_^2^= 0.05, 95% CI [−0.22, 0.84]. In short, the number of eye movements as measured with the HEOG was very small and did not differ across conditions. We conclude that it is unlikely that eye movements spuriously generated the CDA and contralateral alpha suppression differences across reliability conditions.

Although eye movements were very small and were not significantly different across reliability conditions, trial-by-trial differences in eye movements might have affected our EEG signals of interest. In order to evaluate this possibility, we calculated Pearson correlation coefficients across trials, per participant, per condition, between contralateral minus ipsilateral HEOG, and the CDA and contralateral alpha suppression, each averaged throughout the time interval of interest (400-900 ms). If trial-by-trial differences in eye position affected alpha-band power or CDA, then they should be correlated across trials. Contrary to this claim, there was no difference across reliability conditions in Fisher transformed Pearson’s correlations between the contralateral minus ipsilateral HEOG and the CDA, *F*(1, 29) = 0.07, *p* = 0.796, η_p_^2^< 0.01, 95% CI [−0.04, 0.05], or the alpha-band power, *F*(1, 29) = 0.18, *p* = 0.672, η_p_^2^< 0.01, 95% CI [−0.04, 0.06]. Lastly, given that the reliability effects we observed were specific to the early time window for CDA and late time window for contralateral alpha suppression, we repeated the aforementioned trial-wise correlation analysis separately for early (400-514 ms) and late (724-900 ms) time windows at which the aforementioned cluster-based permutation tests showed significant main effects of cue reliability for CDA and contralateral alpha suppression respectively. Again, the average Fisher’s transformed Pearson’s correlation coefficients did not differ across conditions for neither of the time windows (all *p*s > 0.369). Together, these results suggest that the differences in CDA and contralateral alpha suppression across reliability conditions cannot be attributed to differences in eye movements across these conditions.

## Discussion

Selective attention has been claimed to be essential for WM storage^12,14,16,19,69,70^. Yet, results regarding the costs of allocating attention away from WM representations have been conflicting^31–33,37,38,40^. To shed new light on these discrepant results, we used EEG indices of spatial selective attention (i.e. contralateral alpha suppression) and storage (i.e. CDA) within WM when the most task-relevant representation was cued. Critically, following our previous work that shows retro-cue costs are sensitive to the reliability of the cue, we manipulated the proportion of valid to invalid trials of the retro-cues across blocks (80% valid vs. 50% valid).^41^ We replicated our behavioral findings by showing that the effects of retro-cues on a continuous report recall task performance are larger for more reliable cues (i.e. 80% valid).

Importantly, here we show that these behavioral effects have a correlate in the amount of attention paid to the cued item on the one hand, and the dropping of non-cued items on the other. Interestingly, we found that reliability mainly affected the timing and duration of different mechanisms, rather than their amplitude. Specifically, right after the cue, contralateral alpha suppression was equal regardless of cue reliability, suggesting that the cue was used to direct *attention* to the cued representation as it presented useful information both when the cue was 50% valid or 80% valid. However, contralateral alpha suppression then persisted throughout the retention period only for highly reliable cues, while it quickly dropped to baseline for less reliable cues. This indicates that attention was sustained on the cued item when participants could be reasonably sure that it would also be the tested item, while they also reverted to uncued items when there was a decent probability of being tested on those too. As a measure of *storage* we took the CDA, which emerged early in the retention interval after highly reliable cues, consistent with a rapid drop of the uncued item from WM. Later in retention, a CDA also emerged for low-reliability retro-cue condition, suggesting that eventually uncued items were also dropped from WM after less reliable cues. Thus, the time-resolved nature of our EEG measures reveals that the reliability of the retro-cue had dissociable effects on the dynamics of the CDA and contralateral alpha suppression. While for highly reliable cues non-cued items were both unattended and dropped from WM, for less reliable cues non-cued items were initially unattended but kept in WM. These results suggest that attentional selection of an item in WM is not accompanied by the loss of unattended items when there is a relatively high chance that these items could be relevant in the future. Thus, our results support a dissociation between selective attention and storage in WM.

Uncued items were unattended but kept in WM following less reliable retro-cues. To our knowledge, ours is the first study to observe simultaneous neural evidence for prioritization of attended items and active storage of unattended items in WM. This finding is in contrast with the claims that suggest WM storage is a direct reflection of selective attention in WM^14,16,19,69,70^, and supports the view that attentional prioritization of an item is a decision separate from dropping the remaining items^22–27^. This discrepancy between the two bodies of evidence is likely due to differences between the perceived future relevance of unattended items across studies, since studies that observed costs for unattended items used highly reliable cues for which here we show that unattended items were dropped immediately. Yet, also for less reliable cues non-cued items were eventually dropped from WM right before the onset of the test display. There are two explanations for the delayed loss of unattended items for less reliable cues. First, non-cued items might have become more vulnerable to interference from other items in WM due to being initially unattended following the retro-cue. This, in turn might have resulted in the deterioration of non-cued items through the retention interval. This scenario is in line with the evidence that proposes selective attention protects WM items against inter-item interference during storage^25^. Alternatively, non-cued items might have been stored in WM for a longer duration in an attempt to create passive memory traces for these items. Passive memory traces have been claimed to be established over short durations for currently less relevant representations^71–73^. Later, with the anticipation of the test display they might have been deliberately dropped from WM in order to allocate all mnemonic resources to the most relevant item to protect it against perceptual interference by the test display. This strategy would be effective given previous findings that show smaller perceptual interference for smaller memory loads^44–46^. Importantly, although both of these explanations support the protective role of selective attention for storage in WM, they are not against our conclusion that selective attention and storage in WM are distinct constructs, as here we show that an item can be unattended but actively stored in WM at a given time.

At first glance our behavioral findings might seem to contradict our EEG findings. Although the CDA – the WM storage index – at the onset of the memory probe was equal for highly reliable and less reliable retro-cues, errors were larger for highly reliable cues. There are two explanations for this apparent discrepancy that are not mutually exclusive. First, it has been shown that even at short intervals currently less important memory items can be stored as passive memory traces^26,74^. Specifically, recent studies have claimed that unattended items are stored silently, without sustained neural activity, through patterns of synaptic weights^71,75–78^. Therefore, it is possible that, in anticipation of being probed with then, non-cued items in 50% valid condition were stored in passive memory traces that were not reflected in the CDA. In 80% valid condition on the other hand, non-cued items might have been completely forgotten in anticipation of not being probed with them. This explanation is consistent with embedded-component memory models that propose different levels of storage related activity depending on the relevance of stored items^24,27^. Importantly, here we show that unattended items can be stored actively in WM, as suggested by the absence of an emerging CDA following the-retro cue for highly reliable cues. [Although this is an indirect evidence for active WM storage, the alternative – that both attended and unattended items are stored silently, thus generating no lateralized ERP – is extremely given dozens of CDA papers that show sustained CDAs at retention intervals comparable to ours.] Thus, our results suggest that whether an unattended item will be stored actively, stored passively, or forgotten completely depends on its perceived future relevance and can be decided strategically. In line with this claim, studies those that support activity-silent WM used cues that were 100% reliable regarding which item was going to be tested first^71,78^.

Second, although the CDA was equal at the end of the trial across reliability conditions, contralateral alpha suppression was present only in 80% valid condition, which suggests that the cued item was selectively attended only for highly reliable cues prior to the test display. Previous work have shown that selectively attending to a WM item protects it against interference by new perceptual input^44–46^. According to this attentional protection account, it is possible that interference by the test display disrupted uncued items more in 80% valid condition. Thus, allocating attention away from non-cued items might have made them more vulnerable to interference from the test display, resulting in lower behavioral performance when invalidly tested. Importantly, our main conclusion – that unattended items can be kept in or dropped from WM depending on their perceived future task relevance – is based on a dissociation between the CDA and contralateral alpha suppression that provide direct measures of storage and attentional selection within WM prior to any perceptual or response related interference distorts memory representations.

Selective attention to the cued item was sustained till the end of the trial for highly reliable cues, but not for less reliable cues. We propose that the allocation of attention back to non-cued items for less reliable cues reflects an attempt to revive previously unattended items that were being lost. In line with this, we found that larger contralateral alpha suppression predicted smaller errors on a trial-level for 50% reliable valid cue trials. These findings are consistent with recent evidence that suggests weakly encoded representations, which are presumably also the weakly represented ones, are prioritized during WM retention in an attempt to prevent their loss^79^. Attentional reallocation to non-cued items was not observed for highly reliable cues. Given existing evidence that suggests the use of retro-cues is at least partly under strategic control^41,80,81^, we claim that an item that is being lost is attended only when it is highly relevant for the ongoing task. Thus, our results provide evidence for the flexible nature of WM by showing that selective attention can be strategically adjusted based on the perceived future relevance of WM items. In addition to its protective function, selective attention in WM has also been suggested to increase the accessibility of the attended item in a way that it effectively guides behavior in the external world^23,24,82,83^. Thus, we argue that the presence of an additional attentional prioritization mechanism within WM aids flexible behavior in a dynamic world where there are multiple relevant items required for the task at hand whose relative priority changes frequently.

Recently, retro-cue benefits have been claimed to reflect an increase in the accessibility of the cued item in the absence of sustained selective attention^84^. According to this idea, the cued item is first attended and selected in memory. Then, its status is reconfigured in a way to make it more accessible for behavior. After this reconfiguration is complete, sustained attention is not necessary to keep this item in a prioritized accessible state. This theoretical model is in line with the pattern of results in 50% valid blocks of the present study where selective attention to the cued item was not sustained yet there were behavioral benefits for the cued item. It is thus possible that the cued item was reconfigured for accessibility without sustained attention. However, attention was sustained till the end of the trial in 80% valid blocks. Our results therefore show that, while a brief attentional selection might be sufficient for increasing the accessibility of a task-relevant item, highly relevant items are attended in a sustained manner.

The CDA has been traditionally defined as a sustained relative negativity contralateral to the memory items presented on one hemifield of the screen while the other hemifield is ignored. Here, the memory display contained memory items on both hemifield. Thus, we hypothesized that the emergence of a CDA following a retro-cue would mean that the item contralateral to the retro-cue is continued to be stored while item ipsilateral to the retro-cue is dropped^58^. An alternative explanation for the emergence of the CDA is that it reflects a boost for the cued item instead of signaling the loss of the non-cued item.^85^ Given that the CDA reflects storage of the items in the contralateral side^54,68^, an impact on the storage of the cued item should be reflected on the signal contralateral to the *cued item*. However, contrary to this alternative explanation, the retro-cue reliability affected the signal contralateral to the *non-cued item*. This result suggests that the CDA in the present study was a result of dropping the non-cued item instead of a mnemonic boost for the cued item^58,86^. On the other hand, contralateral alpha suppression effect was specific to the contralateral side relative to the cued item instead of the ipsilateral side. This is consistent with this signal reflecting attentional selection instead of distractor suppression^87^ (but see^88^).

It has been argued that the CDA and contralateral alpha suppression are strongly related signals, to the extent that the CDA is an output of an asymmetry in the amplitude of alpha-band oscillations^89^. This would mean that the CDA reflects the same *attentional selection* mechanism that contralateral alpha suppression does, instead of being an index *storage* per se. However, this conclusion was based on an experiment that manipulated neither WM load nor the task demands, thus confounding task relevance and storage. In the present study, by having multiple items with different task-relevance, we show that contralateral alpha suppression and CDA behaved differently across time. Moreover, we did not observe any correlation between the CDA and contralateral alpha suppression across participants (for a similar dissociation, see^52,90^) or within participants on a trial level. Lastly, the retro-cue reliability effect was reflected in differences in the signal ipsilateral to the cued item for the CDA, but contralateral to the cued item for the contralateral alpha suppression. Together, these results strongly suggest that CDA is not simply a reflection of lateral alpha power asymmetries. We are not the first to observe dissociable patterns of lateral alpha power asymmetry and CDA. Previous studies that manipulated task demands observed larger contralateral alpha suppression when a WM item was stored for a more demanding task while the CDA was unchanged^52,91^. Here we extent these findings by showing, across two conditions, both equal CDA and different contralateral alpha power, and also equal contralateral alpha power and different CDAs at different time points within the same dataset, thus provide strong evidence for a dissociation between these two signals. Together, these findings suggest that the CDA reflects storage in WM^57^ and the contralateral alpha suppression reflects allocation of attention within WM^92^ and argue against a recent claim that suggested CDA reflects the current focus of attention instead of storage in WM^93^.

In sum, by manipulating the reliability of retro-cues that indicate which of multiple WM items is most likely to be tested, we show that unattended items were kept longer in WM, but only when there was a relatively high chance that they could later be tested. Thus, we propose that the decision to drop an item from WM is separate than the decision to allocate attention away from it, and that these decisions can be flexibly adjusted based on dynamic changes in the relative importance of WM representations for the task at hand.

## Author Contributions

EG, DvM, MM, and CO designed the study. EG programmed the experiment. EG and KD collected the data. EG analyzed the data. EG, DvM, JF, MM, and CO wrote the manuscript.

## Additional Information

The authors declare no competing interests.

## Acknowledgments

This work was supported by de Nederlandse organisatie voor Wetenschappelijk Onderzoek (The Netherlands Organisation for Scientific Research) to MM and C.N.L.O. (grant number 404-10-004), and European Research Council Consolidator Grant ERC-2013-CoG - 615423 to C.N.L.O. We would like to thank Joram van Driel, Ingmar de Vries and Matti Vuerre for helpful discussions.

